# Recurrent cortical circuits implement concentration-invariant odor coding

**DOI:** 10.1101/294132

**Authors:** Kevin A. Bolding, Kevin M. Franks

## Abstract

Animals rely on olfaction to find food, attract mates and avoid predators. To support these behaviors, animals must reliably identify odors across different odorant concentrations. The neural circuit operations that implement this concentration invariance remain unclear. Here we demonstrate that, despite concentration-dependence in olfactory bulb (OB), representations of odor identity are preserved downstream, in piriform cortex (PCx). The OB cells responding earliest after inhalation drive robust responses in a sparse subset of PCx neurons. Recurrent collateral connections broadcast their activation across PCx, recruiting strong, global feedback inhibition that rapidly suppresses cortical activity for the remainder of the sniff, thereby discounting the impact of slower, concentration-dependent OB inputs. Eliminating recurrent collateral output dramatically amplifies PCx odor responses, renders cortex steeply concentration-dependent, and abolishes concentration-invariant identity decoding.

---

Although the ability to reliably identify objects over a large range of stimulus intensities is a fundamental feature of all sensory systems, the neural circuit mechanisms that implement intensity invariance remain poorly understood. In mammals, odors are detected by individual olfactory sensory neurons expressing just one out of ~1,000 different types of odorant receptor, each of which projects to a specific pair of glomeruli in the olfactory bulb (OB). At low concentrations, odorants selectively bind high-affinity receptors, activating a sparse combination of responsive glomeruli. However, receptor activation, and thus glomerular activation, becomes less specific at higher odorant concentrations (*1-4*), potentially degrading the representation of odor identity. However, psychophysical studies indicate that odors retain their perceptual identities while concentration varies over several orders of magnitude (*5-7*). The olfactory system must therefore transform these concentration-dependent odor responses at early stages of processing into concentration-invariant representations of odor identity.

In OB, odor-responsive mitral/tufted cells fire bursts of action potentials with odor-specific latencies that tile the ~500 ms respiration cycle (*8-11*). However, the mitral/tufted cells that are most sensitive to the odorant, and therefore convey the most specific information about the odor, will always respond earliest, regardless of concentration (*12-15*). Odor information is then diffusely projected from OB to PCx, so that individual PCx neurons can integrate inputs from different combinations of OB glomeruli, producing odor-specific ensembles of neurons distributed across PCx whose concerted activity encodes odor identity (*16-20*). Theoretical studies have suggested that PCx can form concentration-invariant odor representations by selectively responding to the earliest-active OB inputs while ignoring the contribution of inputs arriving later, which may reflect more spurious activation of lower-affinity receptors (*21-23*). However, whether this occurs, and how such a temporal filter would be implemented within PCx, are not known.

## Concentration-invariance emerges in PCx

To address this question, we simultaneously recorded spiking in populations of mitral cells and ipsilateral PCx principal cells in awake, head-fixed mice in response to different odorants presented at multiple concentrations (Fig. 1A). Example odor responses during the first sniff after odor onset are shown for cells in OB and PCx in Fig. 1B. At low concentrations, odors activated small and specific subsets of cells in both OB (Fig. 1B, *left*, Supplementary Fig. 1) and PCx (Fig. 1B, *right*). At higher concentrations, the fraction of activated OB cells increased while the fraction of odor-activated cells in PCx remained constant (Fig. 1B and C). To examine population-level responses, we constructed trial-by-trial response vectors composed of spike counts for each cell in populations of OB and PCx cells (see Methods). We then projected these high-dimensional responses onto three principal components (Fig. 1, D and E). We used mean distances across the first three principal components to quantify the variance of responses to an odor at different concentrations (Δ *conc.*), and compared these to the variance for repeated presentations of each odor at a single concentration (*repeat*) and for responses to different odors (Δ *odor*). Crucially, Δ *conc.* responses and Δ *odor* responses were equally variable in OB (Fig. 1D), whereas Δ *conc.* responses in PCx were significantly less variable than Δ *odor* responses (Fig. 1E). These data indicate that concentration-invariance emerges in PCx from OB input that is highly concentration-dependent. And, given that PCx is driven directly by OB, this result indicates that PCx selectively extracts and preferentially represents the concentration-invariant features of its OB input.

**Fig. 1.**
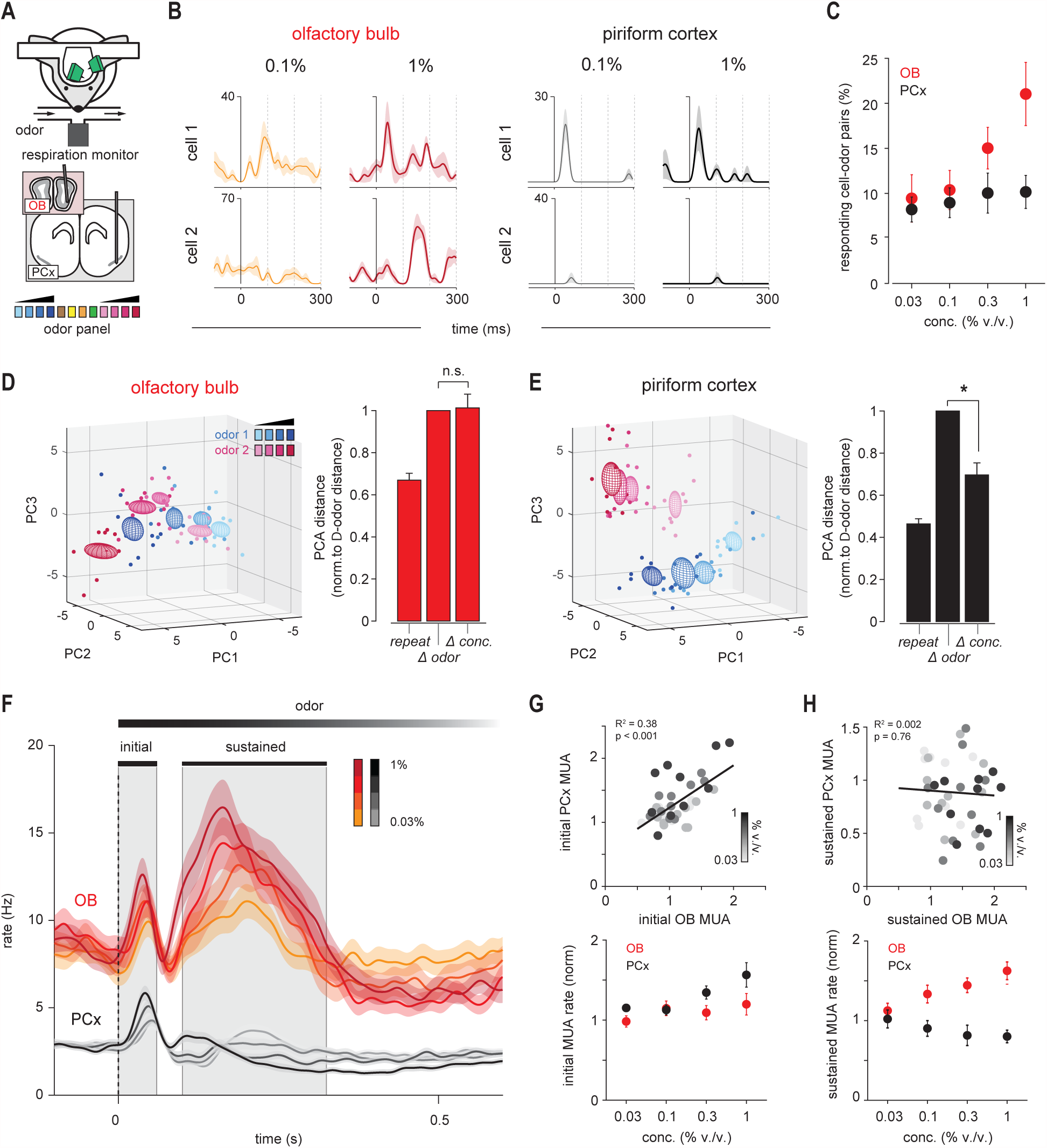
Concentration-invariant odor representations emerge in PCx. (**A**). Experimental schematic. Odor panel included four odors at a single concentration and two odors at four increasing concentrations. (**B**). Example responses from simultaneously recorded pairs of OB (left) or PCx (right) cells to two odors at different concentrations. Responses are aligned to start of inhalation. (**C**) Percent of cells significantly activated by odors of increasing concentration (p < 0.05 rank-sum test, odor vs mineral oil) in OB (red) or PCx (black, n = 5 simultaneous OB-PCx recordings, 2 odors, 4 concentrations). (**D**) Left, PCA representation of OB pseudopopulation response in a 330 ms window after inhalation to ethyl butyrate (blue) and hexanal (magenta) at different concentrations (0.03-1%, different shades). Dots represent responses on individual trials, ellipsoids are mean ± 1 s.d. Right, Relative population response distances in neural activity space projected onto the first three principal components. Distances were computed for each stimulus between trials of the same odor and concentration (repeat, n = 12 stimuli), different odors (Δ odor, n = 12 stimuli), or same odor and different concentration (Δ conc, n = 8 stimuli), and normalized to the average inter-odor distance. OB responses to different concentrations were as dissimilar as responses to different odors (one-sample t-test vs. mean of 1, p = 0.851). (**E**) As in D, but for PCx. PCx responses to different concentrations were more clustered than responses to different odors (p = 0.001). (**F**) Average peristimulus time histograms (PSTHs) for all cell-odor responses at different concentrations (OB, n = 190; PCx, n = 664 cell-odor pairs; mean ± s.e.m.). Gray shading indicates initial (0-60 ms) and sustained (100-300 ms) analysis windows. Dashed line indicates inhalation onset. (**G**) Normalized multiunit activity (MUA) rates during initial phase (n = 5 experiments, 2 odors, 4 concentrations) in OB vs. PCx. Top, each point is the average response of one simultaneously recorded OB-PCx response pair. Shading indicates concentration. Black line is the linear fit. Bottom, Average OB (red) and PCx (black) response across recordings and odors. Multiunit activity was recombined across cells and normalized to baseline activity 1-s before odor. (**H**) As in G but for the sustained phase.

To understand how PCx implements this computation we examined response dynamics over the course of the first sniff. Individual cells in OB and PCx exhibited markedly different dynamics and concentration dependences. Consistent with previous findings (*8-10*), individual OB mitral cells responded with onset latencies that tiled the sniff cycle (Supplementary Fig. 1). This was reflected at the population level as a brief initial increase in spiking followed by a slower and sustained envelope of spiking activity (Fig. 1F, Supplementary Fig. 1). However, in PCx we only observed a transient increase in population spiking followed by sustained suppression, despite continuing input from OB. OB spiking increased systematically with concentration during both the initial and the sustained phases of the response (Fig. 1, G and H). Peak amplitude of the PCx population response did increase, as the responsive ensemble was activated more synchronously at higher concentrations, however the ensembles themselves were largely concentration-invariant (*19*). Notably, the subsequent phase, which was steeply concentration-dependent in OB, was more strongly suppressed for the remainder of the sniff. Thus, PCx may preserve representations of odor identity across concentrations by selectively responding to the earliest OB responses and suppressing its responses to more concentration-dependent inputs that arrive later in the sniff.

## Feedback inhibition truncates PCx odor responses

What is the source of this suppression? Principal neurons in PCx receive inhibitory inputs from two general classes of GABAergic interneurons. Feedforward interneurons reside in layer 1 and only get direct excitatory input from OB (Fig. 2A). These neurons are well positioned to suppress responses to sustained OB input (*24-27*). However, PCx is a highly recurrent circuit in which principal cells extend long-range projections across the cortex, providing excitatory input onto other principal neurons as well as onto feedback interneurons that reside in deep layer 2 and layer 3 (*24, 27-29*). The magnitudes and dynamics of odor-evoked activity in feedforward and feedback inhibitory interneurons in PCx are unknown. We took advantage of the laminar segregation of feedforward and feedback inhibitory interneurons, and used an optical tagging approach to compare odor responses in these two distinct populations of interneurons. We positioned our electrode array in PCx to record from neurons deep or superficial to the large population of glutamatergic principal cells in layer 2 in VGAT-ChR2-GFP mice, in which all GABAergic interneurons express ChR2 (Fig. 2B). Light pulses evoked robust and sustained spiking in ~7% of cells (66/921 cells, n = 15 recordings), consistent with these cells being VGAT+ inhibitory interneurons, while spiking in the majority of the remaining cells was either significantly suppressed (639/921 cells) or unaffected (216/921 cells). We then classified cells as layer 1 feedforward interneurons (FFIs; n = 13/66 VGAT+ neurons) or layer 2/3 feedback interneurons (FBIs; n = 46/66 VGAT+ neurons) according to their DV position relative to the dense population of VGAT-principal cells in layer 2 (Fig. 2, C and D). Seven VGAT+ neurons could not be clearly classified as FFIs or FBIs based on their laminar position and were excluded. Spike waveforms of FBIs were narrower than VGAT-cells and more symmetrical than both VGATand FFIs (Fig. 2D, Supplementary Fig. 2), consistent with a subset of these being fast-spiking interneurons. Spontaneous firing rates in FFIs and FBIs were significantly higher than those in VGAT-cells (Supplementary Fig. 2).

**Fig. 2.**
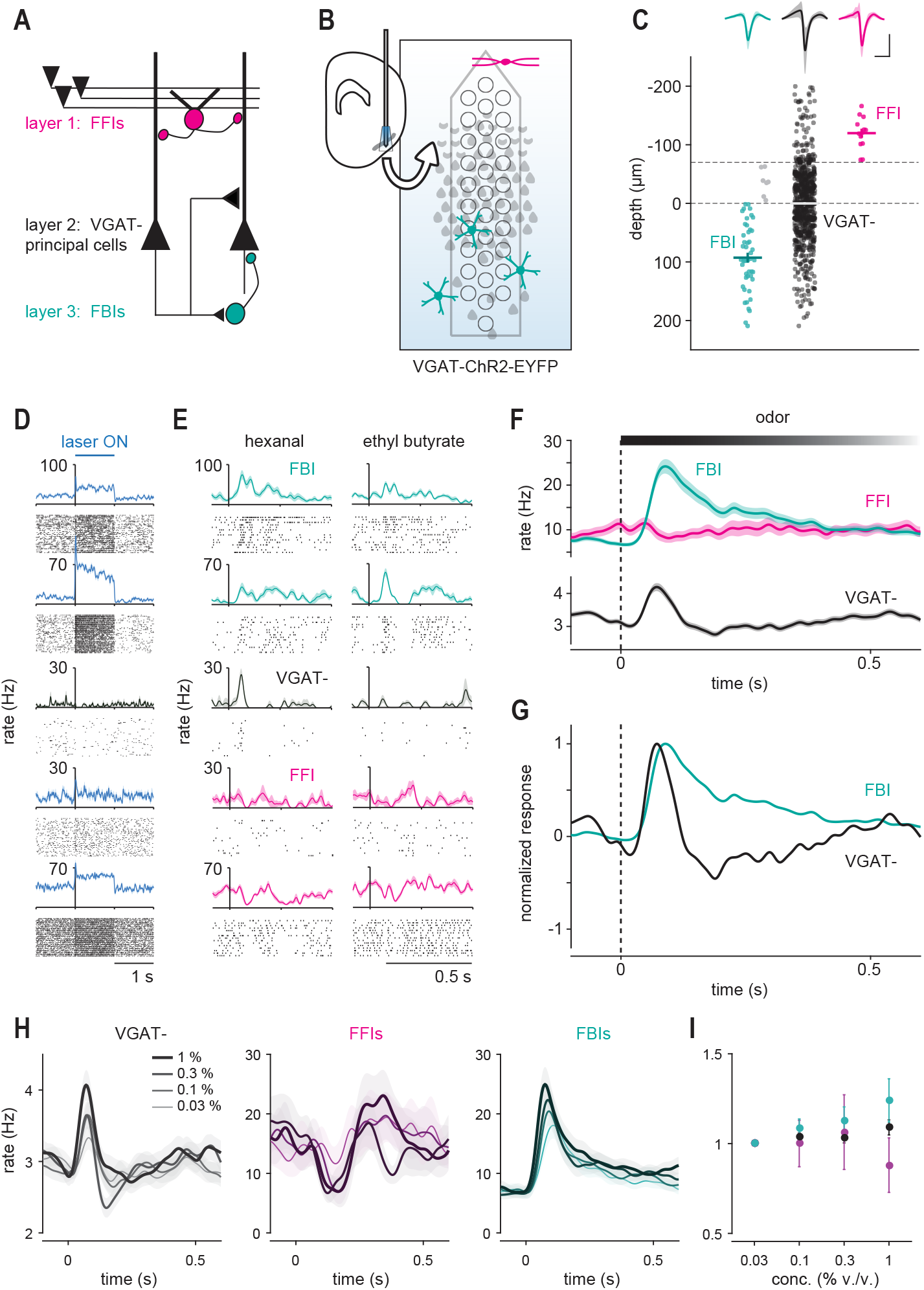
Feedback inhibition shapes cortical odor responses. (**A**) Schematic of PCx circuit: FFIs in layer 1 receive OB input; principal cells in layer 2 provide recurrent excitatory input to other principal cells and to FBIs in layer 3. (**B**) Recording schematic. Light-responsive FFIs and FBIs in VGAT-ChR2 mice are differentiated by their depths relative to VGAT-principal cells in layer 2, which are suppressed. (**C**) FFIs (magenta, n = 13), FBIs (teal, n = 46), and VGAT-(black, n = 855) are classified by light-responsiveness and depth (dashed line). Each point indicates a cell. Thick lines are mean ± s.e.m. Light gray indicates unclassified light-responsive cells. Top, average waveform of each cell-type (mean ± s.e.m., scale bars: 0.5 ms, 0.1 mV). (**D**) Example light responses for one PC (black) and four cells classified as FFIs or FBIs (blue). (**E-F**) Example odor responses (E), and average population PSTHs (F, mean ± s.e.m.) for each cell-type. (**G**) Normalized PSTHs for FBIs and VGAT-cells. (**H**) Average population PSTHs for VGAT-(left), FFIs (middle), and FBIs (right) responding to odors at increasing concentrations. (**I**) Normalized firing rates in response to increasing odor concentrations for each cell type (mean ± s.e.m.).

In response to odors, we observed a large and rapid increase in FBI spiking shortly after inhalation which peaked just as spiking in principal cells was sharply suppressed and which remained elevated for the duration of the sniff (Fig. 2, E to G). By contrast, odor-evoked spiking in FFIs increased slowly and only slightly after inhalation, suggesting that FFIs may provide tonic inhibition driven by spontaneous OB input but do not play a major role in shaping phasic, odor-evoked cortical responses. Furthermore, spiking in FBIs, but not FFIs, increased systematically with concentration (Fig. 2, H and I), consistent with their presumed role in normalizing PCx output. Thus, FBIs appear to play the dominant role in truncating and suppressing odor-evoked activity in PCx. Importantly, because FBIs do not get OB input, but instead are recruited by intracortical recurrent collateral connections, these data indicate that it is PCx activity itself that initiates its subsequent, rapid suppression.

## Recurrent excitation suppresses odor responses

Although intracortical recurrent excitatory circuits are a prominent feature of all sensory cortices, their specific functional and computational roles remain unclear (*30-36*). In PCx, where pyramidal cells receive ~10-fold more recurrent inputs that OB input (*29, 37*), recurrent connections are thought to provide much of the excitatory drive onto odor-responsive cells (*38, 39*). However, because recurrent excitation also recruits FBIs, recurrent circuitry may actually exert a net inhibitory effect on PCx activity. We developed a cortical muting strategy to distinguish between these alternatives. We selectively expressed tetanus toxin light chain (TeLC) in principal cells using cre-dependent AAVs injected into PCx of *emx1-cre* mice, typically infecting ~50% of principal neurons in PCx (Fig. 3, A and B). TeLC expression blocked the release of transmitter from infected neurons (Fig. 3C), without altering their excitability (Fig. 3, D to F) and leaving the rest of the circuitry intact (c.f. (*39*), Supplementary Fig. 3). This strategy allows us to record spiking in a PCx circuit driven directly by OB, without affecting FFI, but eliminating recurrent collateral excitation and consequent recruitment of FBIs. We then recorded bilaterally in PCx after unilateral virus injections. Spontaneous firing rates in TeLC-infected (TeLC-PCx) and contralateral control hemispheres were similar, although population spiking was more strongly coupled to the respiration cycle in both TeLC-PCx and ipsilateral OB, indicating that cortical network activity normally desynchronizes spiking in both PCx and OB (Supplementary Fig. 4).

**Fig. 3.**
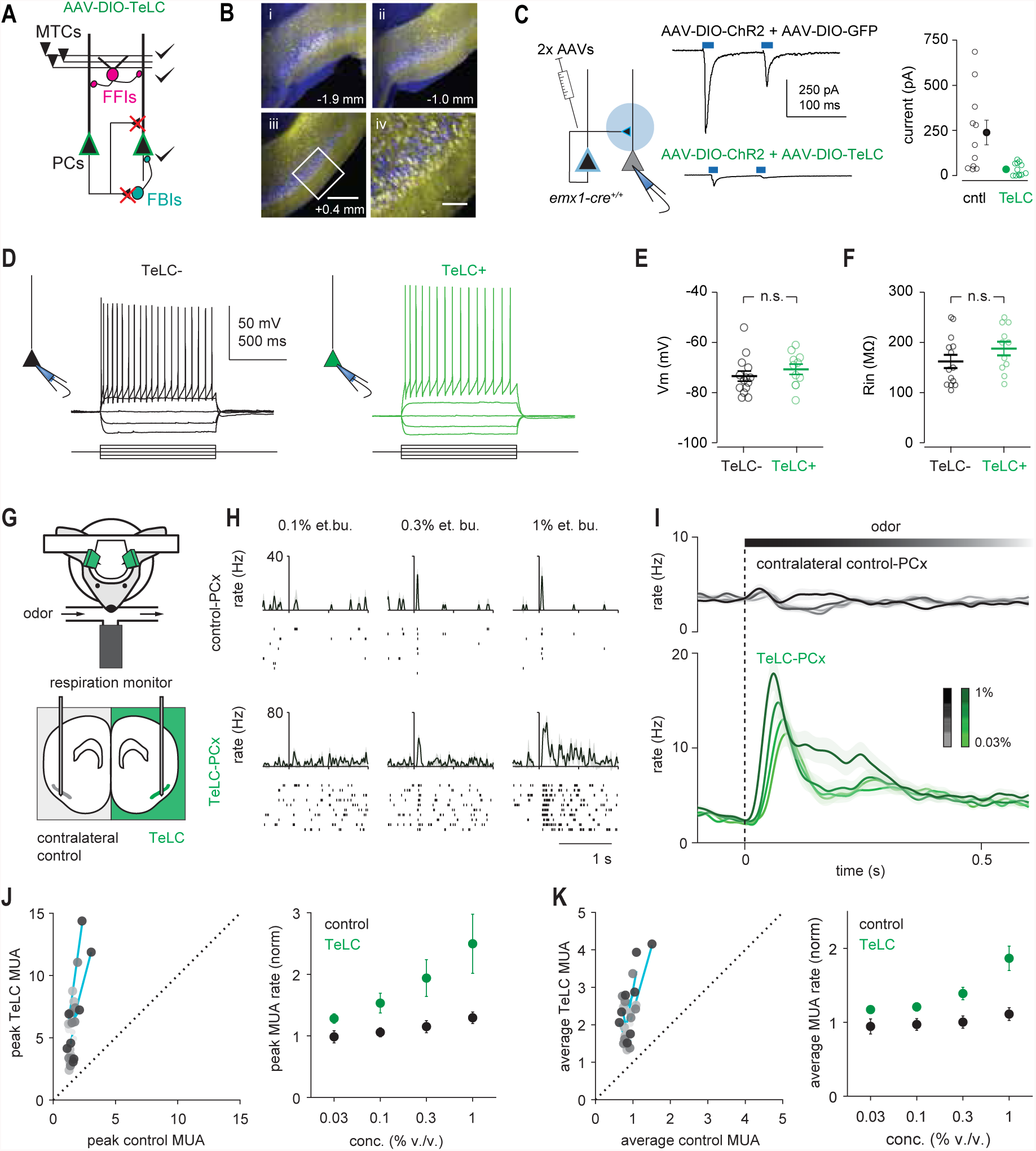
Recurrent circuitry is required to truncate and normalize cortical output. (A)Schematic of circuit changes after selective TeLC expression in PCx principal cells. (B) Extensive infection of layer 2 principal cells across PCx in an example mouse; GFP, yellow; Nissl, blue. Numbers indicate distance from bregma. Figure iv corresponds to inset in iii. Scale bars, 300 μm, 100 μm. (C) Focal co-infection in PCx with ChR2 and either GFP or TeLC-GFP, followed by whole-cell recordings from uninfected cells. Light-evoked synaptic responses are abolished by TeLC. (D) Example recordings from an uninfected (left) and TeLC-infected (right) neuron in the same slice in response to 50 pA current steps. (E-F) Resting membrane potentials (E, TeLC-, 73.4 ± 2.03 mV, n = 14 cells from 3 mice; TeLC+, 70.7 ± 2.01 mV, n = 11 cells from 2 mice; unpaired t-test, p = 0.335) and input resistances (F, TeLC-, 162 ± 13.3 MΩ; TeLC+, 188 ± 13.8 MΩ; p = 0.188) were equivalent. (G) Experimental schematic. Simultaneous bilateral recordings from TeLC-infected and contralateral control hemisphere with odor stimuli. (H-I) Example responses (H) and average population PSTHs (I; mean ± s.e.m., control, n = 450 cell-odor pairs; TeLC, n = 388 cell-odor pairs) (J) Normalized peaks in MUA rates (n = 4 experiments, 2 odors, 4 concentrations). Left, each point is average response of one simultaneously recorded TeLC-Control PCx pair normalized to mineral oil response. Shading indicates concentration. Cyan lines are linear fits for each experiment through all concentrations. Equal concentration-sensitivity in TeLC and control PCx would be indicated by 45° lines. Right, peak responses across recordings and odors. (K) As in G but for average rate over the first 330 ms after inhalation.

Remarkably, despite eliminating much of its excitatory input, odor responses in TeLC-PCx were dramatically enhanced, increasing steeply after inhalation and remaining elevated for the duration of the sniff, (Fig. 3, G to I). Spiking in simultaneously recorded contralateral control hemispheres was truncated shortly after inhalation and suppressed thereafter, as before. Two factors underlie this enhanced population response: first, a given odor activated more and suppressed fewer cells across the population in TeLC-PCx, indicating that recurrent connectivity sparsens cortical odor representations; second, activated responses were larger and of longer duration in TeLC-PCx (Supplementary Fig. 5). We next examined how responses changed across concentrations after eliminating recurrent circuits. Because feedforward inhibition is unaffected by TeLC expression (Supplementary Fig. 3), PCx output should remain stable across concentrations if FFIs control PCx gain. However, we found that response gain was markedly increased in TeLC-PCx, confirming the major role for feedback inhibition in controlling PCx output. Note that gain increased even though odor responses were considerably larger in TeLC-PCx at even the lowest concentrations (Fig. 3, J and K). PCx output remained constant across concentrations in contralateral control hemispheres.

TeLC will block transmitter release from all synapses in infected cells, including centrifugal projections back to OB, as well as to downstream target areas. Centrifugal inputs from PCx contact GABAergic OB neurons that can suppress mitral/tufted cell output (*40-42*), and this process would also be disrupted after TeLC infection. To determine whether the large, prolonged responses observed in TeLC-PCx were simply a consequence of enhanced OB input, we also recorded OB responses ipsi- and contralateral to TeLC-PCx. Indeed, ipsilateral OB responses were markedly larger than contralateral controls and increased more steeply at higher concentrations (Supplementary Fig. 6). However, while the amplitude of the OB response was larger, the time course of the response was unaffected. Thus, centrifugal inputs from PCx play an important role in control the gain of the OB odor response, whereas the marked truncation and sustained suppression after inhalation is implemented within PCx itself.

## PCx responds selectively to the earliest OB inputs

To circumvent the contribution of centrifugal inputs and isolate the intracortical processes that shape PCx odor responses, we used an optogenetic approach to stimulate OB directly. We presented one-second-long light pulses above the dorsal surface of OB of Thy1-ChR2-YFP mice, which express channelrhodopsin-2 (ChR2) in mitral/tufted cells (*43*) (Fig. 4A). For these recordings, we illuminated dorsal OB while recording from mitral cells near the ventrolateral OB surface, providing a lower-bound estimate of the change in OB output. Nevertheless, light pulses elicited an increase in OB spiking that scaled with light intensity and, importantly, remained elevated for the duration of the stimulus (Fig. 4, B to E). This sustained OB activation only produced a large initial peak in PCx population spiking that rapidly returned to baseline for the remainder of the light pulse (Fig. 4C). In fact, although initial peak spike rate in PCx increased steeply at higher light intensities (Fig. 4D), presumably in response to strong and synchronous mitral cell input, sustained population activity was systematically suppressed at higher stimulation intensities (Fig. 4E). Then, as the light pulse ended, the sudden drop in input from OB produced a transient dip in population PCx spiking, which quickly returned to baseline. Thus, PCx dynamically compensates for changes in excitatory drive with rapid recurrent inhibition that balances excitatory input and controls gain to stabilize total cortical output across input intensities. These results are consistent with PCx being an inhibition-stabilized network (*44-47*). These experiments also demonstrate directly that PCx responds robustly to the earliest OB inputs and then suppresses its output in response to OB inputs that arrive later. To reveal the role of recurrent excitation in implementing this transformation, we repeated these experiments in Thy1-ChR2-YFP^+/-^/*emx1-Cre*^+/-^ mice with unilateral TeLC-expression (Fig. 4F). Direct OB stimulation now drove sustained spiking in TeLC-PCx that scaled with intensity (Fig. 4, G to J), similar to what we observed in uninfected, control OB, while recorded responses in contralateral hemispheres were similar to what we observed in uninfected, control PCx. Taken together, our data show that recurrent collateral excitation truncates, suppresses and normalizes PCx odor responses, and therefore has a net-inhibitory effect on cortical activity.

**Fig. 4.**
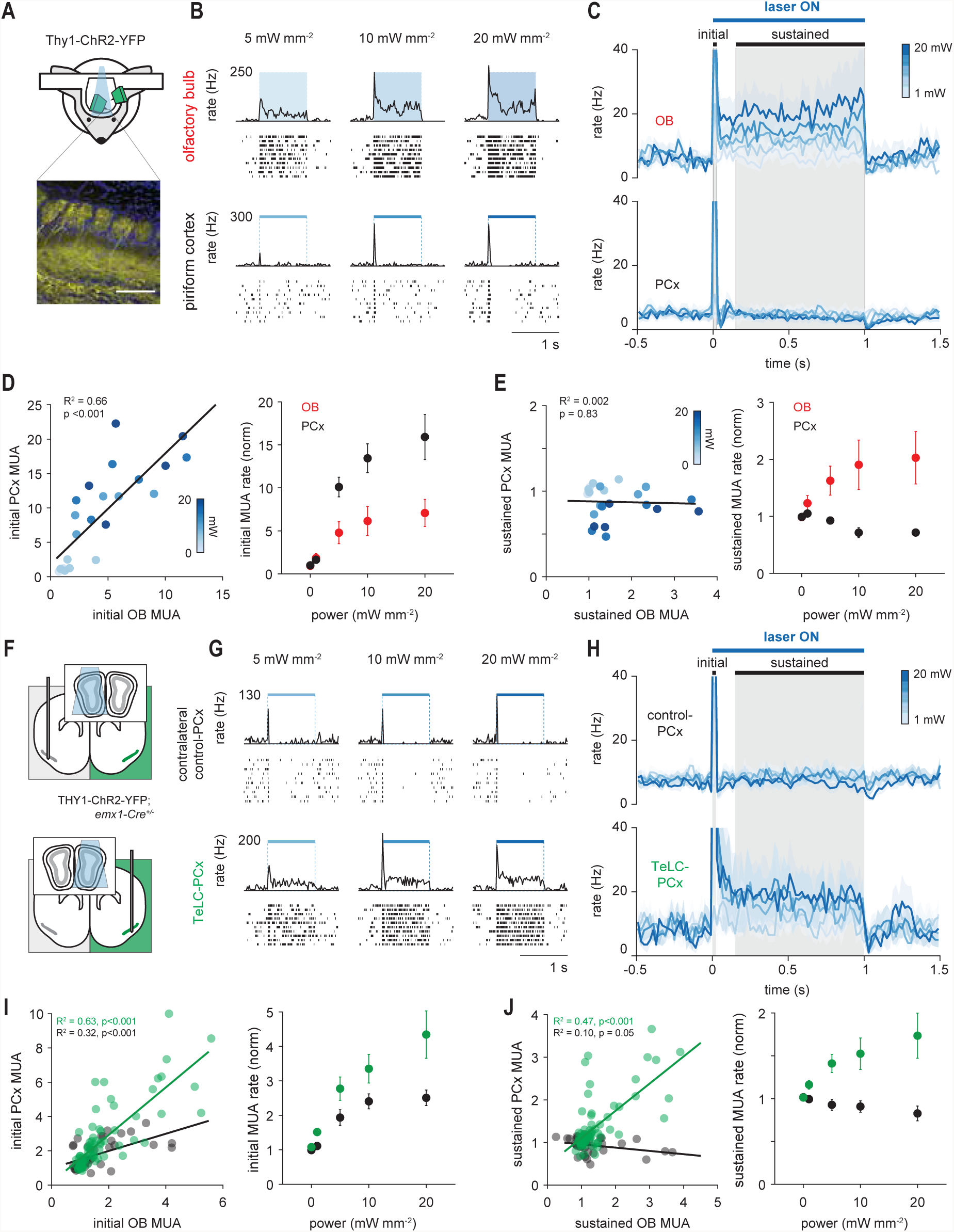
PCx truncates sustained input from OB. (**A** Simultaneous OB-PCx recordings with direct optical OB activation: experimental schematic (top) and ChR2 expression in mitral cells (bottom). Scale bar, 100 μm. (**B**) Responses from example OB (top) and PCx cells (bottom) to 1-second light pulses over OB. (**C**) Average population PSTHs for responses from experiment in B. Gray shading indicates initial and sustained analysis windows. (PCx time constants @ 20 mW; decay from peak: 18.9 ± 2.0 ms; recovery from post-stim. trough: 87.4 ± 46.3 ms; n = 5 population recordings) (**D**) Normalized multiunit activity (MUA) rates during initial phase (n = 5 experiments) in OB vs. PCx. Left, each point is the average response of one simultaneously recorded OB-PCx response pair. Shading indicates light intensity. Black line is the linear fit. Right, Average OB (red) and PCx (black) response across recordings. Multiunit activity was recombined across cells and normalized to baseline activity 1-s before stimulation. (**E**) As in D but for the sustained phase. (**F**) Experimental schematic. Simultaneous OB-PCx recordings from TeLC-infected or contralateral control hemisphere with optical OB activation. (**G**) Example responses from cells to 1-second light pulses over OB. (**H**) Average population PSTHs for responses from experiments in G. (**I**) Normalized peak MUA rates during initial phase (n = 13 TeLC and 8 control experiments) in OB vs. PCx. Left, each point is average response of one simultaneously recorded OB-PCx pair at one concentration. Solid lines are linear fits for TeLC (green) and control (black) data. (**J**) As in I but for sustained rate.

## Recurrent circuitry is required for concentration-invariant decoding

We next asked how eliminating recurrent connectivity alters population odor coding. We performed PCA on single-trial population response vectors from control or TeLC-PCx recordings and calculated the distance between responses in principal component space, as before. In contralateral PCx, Δ *conc*. responses were only slightly more variable than *repeat* responses to a single concentration, and significantly less variable than Δ *odor* responses (Fig. 5A), consistent with results in unperturbed PCx (Fig. 1E). Strikingly, Δ *conc.* responses were much more variable than *repeat* responses and as variable as Δ *odor.* responses in TeLC-PCx (Fig. 5B), equivalent to what we observed in OB under control conditions (Fig. 1E). Thus, recurrent circuits are responsible for the emergence of concentration-invariance in PCx.

**Fig. 5.**
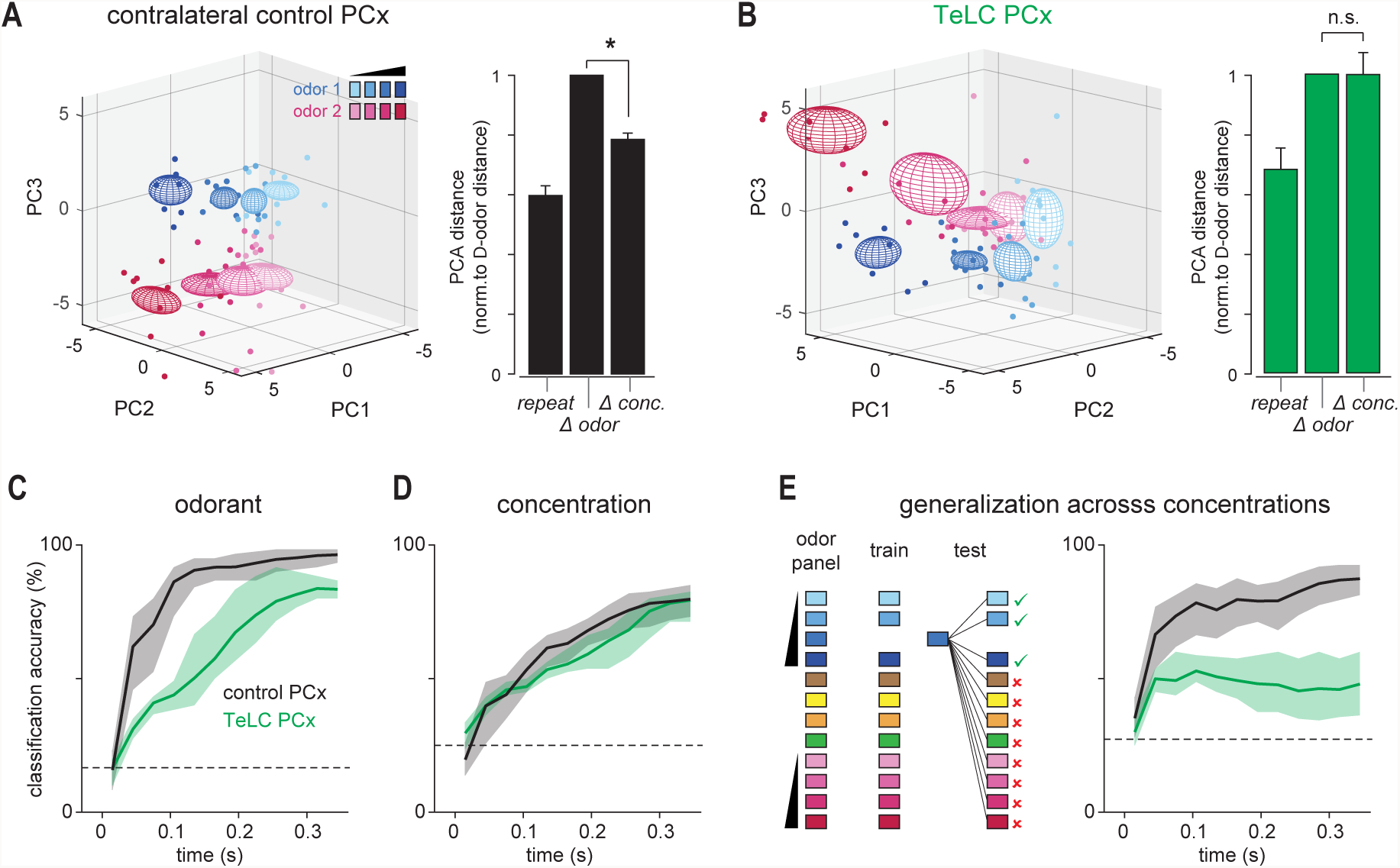
Recurrent circuits implement concentration-invariant decoding. (**A**) Left, PCA representation of pseudopopulation response to ethyl butyrate (blue) and hexanal (magenta) at different concentrations (0.03-1%, different shades). Dots represent responses on individual trials, ellipsoids are mean ± 1 s.d. Right, Mean distance between population responses in PCA space normalized to odor responses. Δ conc. responses (n = 8 stimuli) were more similar than Δ odor responses (n = 12 stimuli) in the control PCx (one-sample t-test vs. mean of 1, p = 2.03e-5). (**B**) As in A, but for TeLC PCx. Δ conc. responses were no more similar than Δ odor responses (p = 0.985). (**C**) Linear classifier performance for odorant decoding (choose 1/6 odors) using TeLC-infected (green) or contralateral control (black) PCx pseudopopulations. Classifier was trained and tested on spike counts in 20-ms bins in an expanding time window starting at odor inhalation. Pseudopopulation size in both conditions was held at 180 cells. Mean ± 95% confidence intervals from 200 permutations. (**D**) Same as C for classification of different concentrations of the same odorant (choose 1/4 dilutions). (**E**) Accuracy for generalization task in which classifier is trained and tested on different concentrations of odors. Loss of recurrent circuits severely impairs odor identity recognition across concentrations.

Finally, we asked how and when odor information becomes available to a downstream observer, and how this is altered when recurrent circuitry is removed. We trained and tested a linear classifier on three decoding tasks: classifying responses to different odorants, classifying responses to a single odorant at different concentrations, and generalizing for odor identity across odorant concentrations (Fig. 5, C to E). Input to the classifier consisted of spike counts for each neuron in an expanding series of 20 ms bins starting with inhalation onset (*9*). Decoding accuracy using responses recorded from the contralateral control hemisphere increased rapidly after inhalation and remained elevated for the duration of the sniff when classifying responses to different odorants or when generalizing for odor identity; concentration decoding was delayed, increased more slowly, and was less accurate over the full sniff. These data indicate that the earliest PCx responses encode largely concentration-invariant representations of odor identity while odor concentration information arrives later, and are consistent with our previous findings in control mice (*19*).

Eliminating recurrent circuitry effectively abolished the ability to generalize for odor identity; decoding accuracy increased slightly immediately after inhalation, but there was no subsequent improvement and, if anything, a small decrease in decoding accuracy as the sniff progressed (Fig. 5E). This generalization result contrasts with performance on classifying responses to different odorants which, while impaired in TeLC-PCx compared to controls, improved slowly but steadily over the sniff (Fig. 5C). Concentration decoding performance was equivalent in control and TeLC-PCx (Fig. 5D). Taken together, we interpret these results to indicate that cortical responses to different odors remain somewhat distinct across the entire sniff but that only the earliest PCx responses convey concentration-invariant, identity-specific information. In control hemispheres, the relative impact of these early cortical responses is amplified by broadcasting their activity across PCx via long-range recurrent collateral connections that effectively recruit feedback inhibitory neurons and, consequently, rapidly and globally suppress subsequent cortical activity for the duration of the sniff. However, when recurrent output is blocked the early responses cannot suppress consequent activity and so PCx continues to be driven by OB input that conveys ever-less identity-specific and more concentration-dependent information as the sniff progresses (*15*). Ultimately, we want to know whether disrupting this circuitry abolishes concentration-invariant odor perception. However, expressing TeLC in PCx principal neurons blocks transmitter release from all their synapses, including PCx outputs, which precludes behavioral testing. Moreover, direct silencing of feedback interneurons will result in regenerative, epileptogenic activity in the highly recurrent circuit. Therefore, development of optogenetic or chemogenetic effectors that can be efficiently targeted to defined subsets of synapses will be required to reveal the behavioral consequences of disrupting recurrent connectivity.

We have revealed an essential role for recurrent feedback inhibition in preserving representations of odor identity across odorant concentrations. The combination of recurrent excitation and feedback inhibition implements a “temporal winner-take-all” filter to extract and selectively represent the most concentration-invariant features of the odor stimulus. This process emphasizes the earliest and most odor-specific inputs to PCx. Similar types of “first-spike” coding strategies have been identified in the visual (*48, 49*), somatosensory (*50, 51*) and auditory systems (*52*). Sensory representations are topographically ordered in these neocortical sensory areas, which allows local, surround inhibition to implement this temporal filter (*53, 54*). However, odor ensembles are distributed across PCx and lack any discernable topographic organization (*17, 20*). Consequently, diffuse, long-range recurrent collateral projections that recruit strong feedback inhibition ensure that recurrent inhibition is, effectively, global in PCx (*29*). This global inhibition truncates activity, sparsens responses, controls cortical gain, and supports concentration-invariant representations of odor identity. Thus, although recurrent circuitry in PCx is typically thought to provide the excitatory substrate for odor learning, memory and olfactory pattern completion (*55, 56*), we find that recurrent excitation has a net-inhibitory impact on cortical activity. In fact, strong and global feedback inhibition that sparsens and normalizes output has been identified at the equivalent stage of processing in invertebrate olfactory systems, however this is implemented by a single, globally-connected interneuron (*57, 58*). Interestingly, the highly-recurrent CA3 region of hippocampus exhibits a similar pattern of long-range recurrent collateral connectivity (*59*). Thus, recurrent excitation that is dominated by rapid, global feedback inhibition may reflect a canonical circuit motif for temporally filtering representations in associative cortex and related structures.

## Methods

### Subjects

All experimental protocols were approved by Duke University Institutional Animal Care and Use Committee. The methods for head-fixation, data acquisition, electrode placement, stimulus delivery, and analysis of single-unit and population odor responses are adapted from those described in detail previously (19). A portion of the data reported here (5 of 13 simultaneous OB and PCx recordings) were also described in that previous report. Mice were singly-housed on a normal light-dark cycle. For simultaneous OB/PCx recordings and Cre-dependent TeLC expression experiments, mice were adult (>P60, 20-24 g) offspring of Emx1-cre (+/+) breeding pairs obtained from The Jackson Laboratory (005628). Optogenetic experiments used adult Thy1-ChR2-YFP (+/+), line 18 (Jackson Laboratory, 007612) and VGAT-ChR2-YFP (+/-), line 8 (Jackson Laboratory, 014548). Adult offspring of Emx1-cre (+/+) mice crossed with Thy1-ChR2-YFP (+/+) mice were used for combined optogenetics and TeLC expression.

### Adeno-associated viral vectors

All viruses were obtained from the vector core at the University of North Carolina-Chapel Hill (UNC Vector Core). AAV5-CBA-DIO-TeLC-GFP, AAV5-CBA-DIO-GFP, AAV5-ef1a-DIO-ChR2-EYFP were used for in vitro slice physiology experiments. For in vivo experiments, AAV5-DIO-TeLC-GFP was expressed either under control of a CBA (4/5) or synapsin (1/5) promoter and results were pooled. TeLC expression throughout PCx was achieved using 500 nL injections at three stereotaxic coordinates (AP, ML, DV: +1.8, 2.7, 3.85; +0.5, 3.5, 3.8; -1.5, 3.9, 4.2; DV measured from brain surface). Recordings were made ~14 days post-injection. To confirm widespread expression of TeLC in PCx, after all recordings, mice were perfused with 4% PFA, coronal sections were taken through the A-P extent of PCx, and slices were labeled with anti-GFP (1:500, Cell Signaling Technology).

### In vitro electrophysiology and Analysis

For experiments examining viability and excitability in TeLC-infected neurons (Supplementary Figs 3, A-C, G-I), viruses were injected as described above. For experiments validating that transmitter release was blocked in TeLC-infected neurons and OB inputs were unaffected (Fig. 3B and Supplementary Figs. 3D-F) a single injection containing a cocktail of 150 nL AAV-EF1 a-DIO-ChR2-EYFP and either 150 nL AAV-EF1 a-DIO-ChR2-GFP or 150 nL AAV-CAG-DIO-GFP-TeLC were injected at a single site in anterior PCx.

Fifteen ± 2 days after virus injection, mice were anesthetized with isofluorane and decapitated. The cortex was quickly removed in ice-cold artificial CSF (aCSF). Parasagittal brain slices (300 μm) were cut using a vibrating microtome (Leica) in a solution containing (in mM): 10 NaCl, 2.5 KCl, 0.5 CaCl2, 7 MgSO4, 1.25 NaH2PO4, 25 NaHCO3, 10 glucose, and 195 sucrose, equilibrated with 95% O2 and 5% CO2. Slices were incubated at 34°C for 30 min in aCSF containing: 125 mM NaCl, 2.5 mM KCl, 1.25 mM NaH2PO4, 25 mM NaHCO3, 25 mM glucose, 2 mM CaCl2, 1 mM MgCl2, 2 NaPyruvate. Slices were then maintained at room temperature until they were transferred to a recording chamber on an upright microscope (Olympus) equipped with a 40x objective.

For current clamp recordings, patch electrodes (3-6 MΩ) contained: 130 Kmethylsulfnoate, 5 mM NaCl, 10 HEPES, 12 phosphocreatine, 3 MgATP, 0.2 NaGTP, 0.1 EGTA, 0.05 AlexaFluor 594 cadaverine. For voltage-clamp experiments, electrodes contained: 130 D-Gluconic acid, 130 CsOH, 5 mM NaCl, 10 HEPES, 12 phosphocreatine, 3 MgATP, 0.2 NaGTP, 10 EGTA, 0.05 AlexaFluor 594 cadaverine. Voltage- and current-clamp responses were recorded with a Multiclamp 700B amplifier, filtered at 2-4 kHz, and digitized at 10 kHz (Digidata 1440). Series resistance was typically ~10 MΩ, always <20 MΩ, and was compensated at 80%–95%. The bridge was balanced using the automated Multiclamp function in current clamp recordings. Data were collected and analyzed off-line using AxographX and IGOR Pro (Wavemetrics). Junction potentials were not corrected.

Recordings targeted pyramidal cells, which were visualized (CoolLED) to ensure that cells had pyramidal cell morphologies. For current clamp recordings to examine viability and excitability, TeLC- or GFP-infected neurons were targeted using 470 nm light (CoolLED). In current clamp recordings, a series of 1 s. current pulses were stepped in 50 pA increments.

To examine synaptic properties, we first verified that fluorescent cells exhibited large photocurrents in both ChR2-YFP/GFP- and ChR2-YFP/GFP-TeLC-injected slices (not shown). We then recorded in voltage-clamp from uninfected cells adjacent to the infection site. Cells were held at either -70 mV or +5 mV to isolate excitatory or inhibitory synaptic currents, respectively. Brief (1 ms) 470 nm pulses were delivered through the objective every 10 seconds to activate ChR2+ axon terminals. A concentric bipolar electrode in the lateral olfactory tract was used to activate synaptic inputs from OB (Supplementary Fig. 3, D-F). The bipolar electrode was placed at the layer 2/3 border 226 ± 17 mm from the recorded cell to examine feedback inhibition (Supplementary Fig. 3, G-I). All viruses were obtained from the UNC Viral Vector Core facility. NBQX, D-APV and gabazine were acquired from Tocris.

### Head-fixation

Mice were habituated to head-fixation and tube restraint for 15-30 minutes on each of the two days prior to experiments. The head post was held in place by two clamps attached to ThorLabs posts. A hinged 50 ml Falcon tube on top of a heating pad (FHC) supported and restrained the body in the head-fixed apparatus.

### Data acquisition

Electrophysiological signals were acquired with 32-site polytrode acute probes (A1x32-Poly3-5mm-25s-177, Neuronexus) through an A32-OM32 adaptor (Neuronexus) connected to a Cereplex digital headstage (Blackrock Microsystems). A fiber-attached polytrode probe (A1x32-Poly3-5mm-25s-177-OA32LP, Neuronexus) was used for recordings from optogenetically identified GABAergic cells. Unfiltered signals were digitized at 30 kHz at the headstage and recorded by a Cerebus multichannel data acquisition system (BlackRock Microsystems). Experimental events and respiration signals were acquired at 2 kHz by analog inputs of the Cerebus system. Respiration was monitored with a microbridge mass airflow sensor (Honeywell AWM3300V) positioned directly opposite the animal’s nose. Negative airflow corresponds to inhalation and negative changes in the voltage of the sensor output.

### Electrode and optic fiber placement

The recording probe was positioned in the anterior piriform cortex using a Patchstar Micromanipulator (Scientifica). For piriform cortex recordings, and the probe was positioned at 1.32 mm anterior and 3.8 mm lateral from bregma. Recordings were targeted 3.5-4 mm ventral from the brain surface at this position with adjustment according to the local field potential (LFP) and spiking activity monitored online. Electrode sites on the polytrode span 275 μm along the dorsal-ventral axis. The probe was lowered until a band of intense spiking activity covering 30-40% of electrode sites near the correct ventral coordinate was observed, reflecting the densely packed layer II of piriform cortex. For standard recordings, the probe was lowered to concentrate this activity at the center of the DV axis of the probe. For deep or superficial recordings, the probe was targeted such that strong activity was at the most ventral or most dorsal part of the probe respectively. For simultaneous ipsilateral olfactory bulb recordings, a micromanipulator holding the recording probe was set to a 10-degree angle in the coronal plane, targeting the ventrolateral mitral cell layer. The probe was initially positioned above the center of the olfactory bulb (4.85 AP, 0.6 ML) and then lowered along this angle through the dorsal mitral cell and granule layers until encountering a dense band of high-frequency activity signifying the targeted mitral cell layer, typically between 1.5 and 2.5 mm from the bulb surface. For experiments driving OB cells in Thy1-ChR2-YFP mice, an optic fiber was positioned <500 μm above the dorsal surface of the bulb.

### Spike sorting and waveform characteristics

Individual units were isolated using Spyking-Circus (https://github.com/spyking-circus)(60). Clusters with >1% of ISIs violating the refractory period (< 2 ms) or appearing otherwise contaminated were manually removed from the dataset. This criterion was relaxed to 2% in Thy1-ChR2-YFP recordings because these were short (<15 minutes) and had poorer overall sorting quality, and these results do not depend on unit isolation, but rather total population spiking activity. Pairs of units with similar waveforms and coordinated refractory periods in the cross-correlogram were combined into single clusters. Extracellular waveform features were characterized according to standard measures: peak-to-trough time and ratio and peak amplitude asymmetry (61). Unit position with respect to electrode sites was characterized as the average of all electrode site positions weighted by the wave amplitude on each electrode.

### Spontaneous activity and respiration-locking

Spontaneous activity was assessed during inter-trial intervals at least 4 seconds after stimulus offset and 1 second preceding stimulus. The relationship of each unit’s spiking to the ongoing respiratory oscillation was quantified using both phase concentration (κ) (62) and pairwise phase consistency (PPC) (63). Each spike was assigned a phase by interpolation between inhalation (0 degrees) and exhalation (180 degrees). Each spike was then treated as a unit vector and PPC was taken as the average of the dot products of all pairs of spikes.

### Individual and average cell-odor responses

We computed smoothed kernel density functions (KDF) with a 10 ms Gaussian kernel (using the psth routine from the Chronux toolbox: www.chronux.org)(64) to visualize trial-averaged firing rates as a function of time from inhalation onset and to define response latencies for each cell-odor pair. Multi-unit activity or population responses were constructed by averaging these KDFs across all cells and odors. Peak latency was defined as the maximum of the KDF within a 500-ms response window following inhalation. Response duration was the full-width at half-maximum of this peak.

### Identifying VGAT+ interneurons

To assess odor responses in identified interneurons, 1-s light pulses were delivered just above the recording sites using a fiber-attached probe. Twenty pulses were delivered both before and after presentation of the full odor stimulation series. Cells were labeled as laser-responsive using a Wilcoxon rank-sum test comparing firing rates in the 1-s prior to and during laser stimulation.

### Sparseness

Lifetime and population sparseness were calculated as described previously (19, 65).

### Principal components analysis

Principal components were computed from pseudo-population response vectors using the Dimensionality Reduction Toolbox (https://lvdmaaten.github.io/drtoolbox/). Response were spike counts over the first 330 ms after inhalation on each trial for each cell. Responses were combined across all cells in TeLC or control conditions to form pseudo-population response vectors. To compute PC distance, 3-dimensional Euclidean distances were computed for each trial pair and the average trial-pair distance was computed for each stimulus. For example, the “same-odor, different concentration” distance for 1% ethyl butyrate is an average of ten 1% trials’ distances from thirty trials of three other concentrations (300 distances). Summary statistics were computed on these average trial-pair distances.

### Population decoding analysis

Odor classification accuracy based on population responses was measured using a Euclidean distance classifier with Leave-One-Out cross-validation. Responses to four distinct monomolecular odorants presented at 0.3% v/v and two more odorants presented in a concentration series at 0.03%, 0.1%, 0.3% and 1% v/v were used as the training and testing data. For generalization tasks, one concentration was left out during training and testing and the classifier prediction was recoded as ‘correct’ if the predicted odor was of the same identity as the presented odor. The feature vectors were spike counts in concatenated sets of 20 ms bins over the first 340 ms following inhalation.

### Statistics

Statistics were computed in MATLAB. Paired t-tests were used when comparing the same animals, cells, or cell-odor pairs across states. Unpaired t-tests and two-sample Kolmogorov-Smirnov tests were used when comparing properties for distinct cell-odor pairs. Sample sizes were large such that t-tests were robust to non-normality. Results were equivalent with non-parametric tests. No formal a priori sample size calculation was performed, but our sample sizes are similar to those used in previous studies.

## Acknowledgements

We thank P. Lee and M. Dalgetty for technical assistance; and A. Fleischmann, S. Lisberger, R. Mooney, M. Scanziani, A. Schaefer, M. Tadross and F. Wang for reading early versions of the manuscript. This work was supported by grants from NIDCD (DC009839 and DC015525) and the Edward Mallinckrodt Jr. Foundation.

